# Pediatric Cancer Variant Pathogenicity Information Exchange (PeCanPIE): A Cloud-based Platform for Curating and Classifying Germline Variants

**DOI:** 10.1101/340901

**Authors:** Michael N. Edmonson, Aman N. Patel, Dale J. Hedges, Zhaoming Wang, Evadnie Rampersaud, Chimene A. Kesserwan, Xin Zhou, Yanling Liu, Scott Newman, Michael C. Rusch, Clay L. McLeod, Mark R. Wilkinson, Stephen V. Rice, Jared B. Becksfort, Kim E. Nichols, Leslie L. Robison, James R. Downing, Jinghui Zhang

## Abstract

Variant interpretation in the era of next-generation sequencing (NGS) is challenging. While many resources and guidelines are available to assist with this task, few integrated end-to-end tools exist. Here we present “PeCanPIE” – the Pediatric Cancer Variant Pathogenicity Information Exchange, a web- and cloud-based platform for annotation, identification, and classification of variations in known or putative disease genes. Starting from a set of variants in Variant Call Format (VCF), variants are annotated, ranked by putative pathogenicity, and presented for formal classification using a decision-support interface based on published guidelines from the American College of Medical Genetics and Genomics (ACMG). The system can accept files containing millions of variants and handle single-nucleotide variants (SNVs), simple insertions/deletions (indels), multiple-nucleotide variants (MNVs), and complex substitutions. PeCanPIE has been applied to classify variant pathogenicity in cancer predisposition genes in two large-scale investigations involving >4,000 pediatric cancer patients, and serves as a repository for the expert-reviewed results. While PeCanPIE’s web-based interface was designed to be accessible to non-bioinformaticians, its back end pipelines may also be run independently on the cloud, facilitating direct integration and broader adoption. PeCanPIE is publicly available and free for research use.

## Introduction

Next-generation sequencing (NGS) has quickly become a mainstay for genetic variation studies in many research and clinical genomics laboratories. However, the sheer abundance of data produced for a single individual means that complex and often tedious data processing and curation are required to identify potentially disease-causing mutations. The process is simultaneously burdened by the volume of novel variants, many of which have scarce information available, and the diverse, distributed nature of existing variant information resources. Variant annotation tools have been developed to assist with several aspects of this work, which can add coding and noncoding prediction annotations and population-specific allele frequencies, as well as provide filtering options for variant prioritization (Wang et al. 2010; Cingolani et al. 2012; Ng et al. 2009; McLaren et al. 2016). Likewise, variant curation tools supporting classification for clinical pathogenicity following the ACMG guidelines (Richards et al. 2015) have also been developed (Patel et al. 2017). While each resource offers valuable information to help researchers classify variant pathogenicity, integrated platforms are needed to provide support for all steps of the process, and streamline analysis of the thousands to millions of variants generated by NGS-based platforms. With these goals in mind, we developed “PeCanPIE” – the Pediatric Cancer Variant Pathogenicity Information Exchange – a cloud-based portal that provides an end-to-end workflow, beginning with a set of variants in VCF (Danecek et al. 2011) and ending with formal ACMG classification. PeCanPIE offers three key functions: 1) automated annotation, classification, and triage via our MedalCeremony pipeline (Zhang et al. 2015); 2) an interactive variant page and visualization tools to support expert curation and committee review; and 3) a reference database of expert-reviewed germline cancer-predisposing mutations.

## Results

### Process overview

As outlined in Fig. 1A, PeCanPIE launches with an interface for uploading a VCF file, which is then filtered to a set of disease-related genes (Methods, Table S1); users may alternatively specify their own list of genes of interest. Variants are next assigned gene and protein annotations and filtered by functional class and population frequency derived from the Exome Aggregation Consortium (ExAC) database (Lek et al. 2016). To ensure that pathogenic germline variants in cancer patients are retained, PeCanPIE uses the distribution of ExAC that excludes patient samples from The Cancer Genome Atlas (TCGA) (McLendon et al. 2008). The remaining variants are stratified into three tiers (gold, silver, and bronze) as an indication of potential pathogenicity computed by our MedalCeremony pipeline. Finally, each “medaled” variant is linked to a standalone page featuring an interface to support semi-automated pathogenicity classification using ACMG guidelines. Two examples in Fig. 1 demonstrate the classification process using VCF files generated from whole-exome sequencing (WES) of an acute lymphoblastic leukemia (ALL) patient (Moriyama et al. 2015) (Fig. 1B) and whole-genome sequencing (WGS) from the Genome in a Bottle (GiaB) project (Zook et al. 2014) (Fig. 1C), respectively. Only 14 of the 63,109 variants from the WES data and 17 of the approximately 4 million variants from the WGS data required expert review, which resulted in 1 and 0 pathogenic/likely pathogenic (P/LP) variants, respectively.

**Figure 1.**
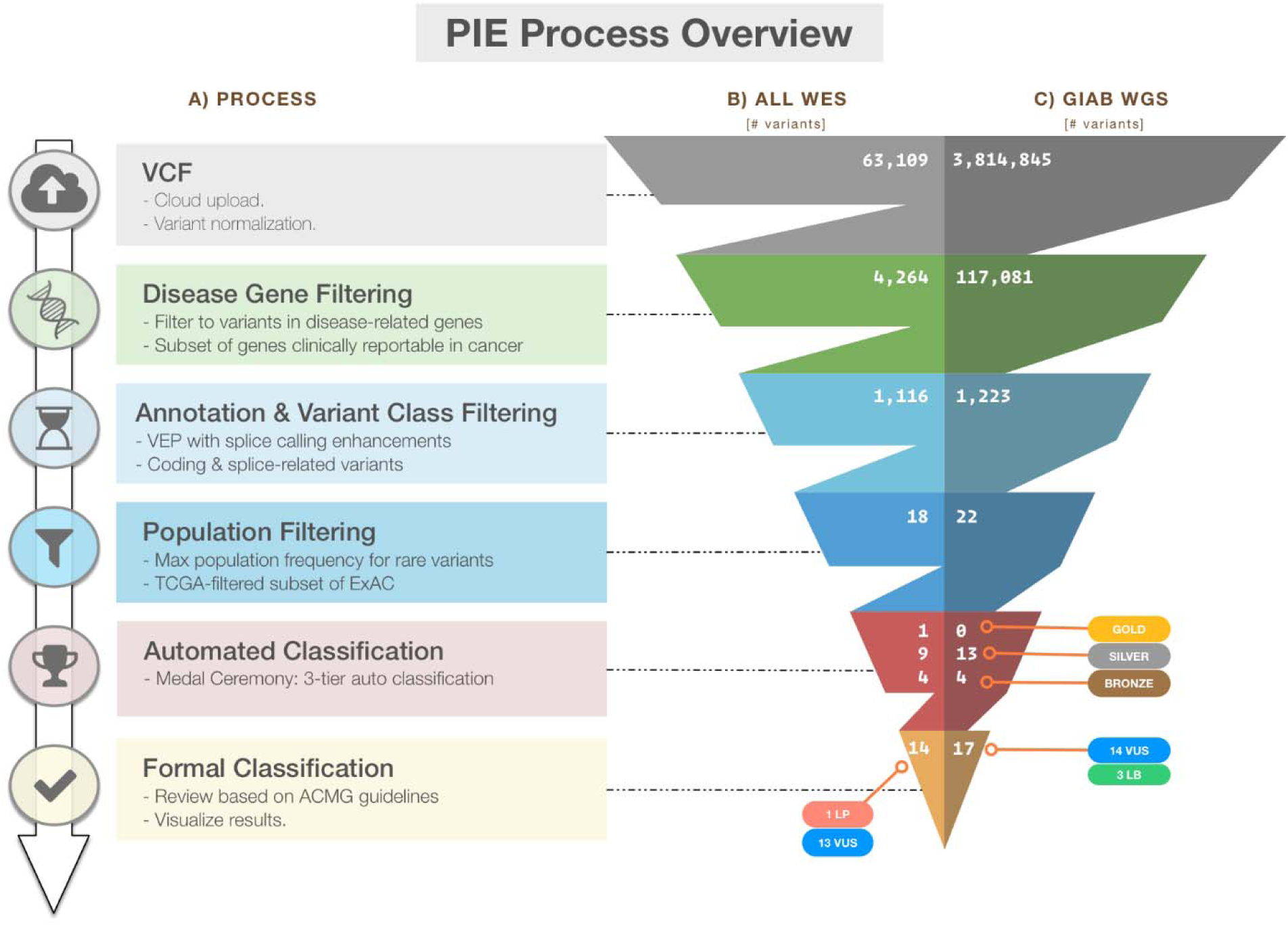
Overview of variant classification using PeCanPIE. (A) Overview of processing steps from VCF through ACMG-based classification. Variant counts at each processing step for (B) whole-exome sequencing data generated from a germline sample of a patient with acute lymphoblastic leukemia (ALL), SJNORM015857_G1 (Methods) and (C) whole-genome sequencing data generated from Genome in a Bottle normal sample NA12878_HG001 (Methods).

### Automated classification by the MedalCeremony pipeline

Automated classification of variant pathogenicity implemented in the MedalCeremony pipeline classifies variants having a population frequency no higher than 0.001 (or a user-defined cutoff) in the ExAC database. Additional annotations are incorporated to aid with the classification process: 1) COSMIC (Forbes et al. 2008) hits; 2) functional annotations from dbNSFP (Liu et al. 2013) (protein domain and damage prediction algorithm calls); and 3) allele frequencies in the NHLBI GO Exome Sequencing Project (ESP), the Thousand Genomes Project (Auton et al. 2015), ExAC, and the Pediatric Cancer Genome Project (PCGP) (Downing et al. 2012).

An overview of the gold, silver, and bronze classification scheme implemented in MedalCeremony is shown in Fig. 2. Gold medals are assigned to truncating variants (including splice variants) in tumor suppressor genes (Zhao et al. 2016; Chakravarty et al. 2017), matches to highly-curated databases (IARC TP53 (Bouaoun et al. 2016), ClinVar expert-panel-reviewed pathogenic (P) or likely pathogenic (LP) variants, ASU TERT (Podlevsky et al. 2007), ARUP RET (Margraf et al. 2009), NHGRI Breast Cancer Information Core (Szabo et al. 2000), somatic mutation hotspots in COSMIC (observed in ≥10 tumors after removal of hypermutators) and PCGP, and St. Jude committee-reviewed germline P/LP variants. Silver medals are assigned to in-frame indels, truncation events in non-tumor-suppressor genes, variants predicted damaging by *in silico* algorithms, and matches to additional databases (ClinVar non-expert-panel P/LP, BRCA Share (Béroud et al. 2016), LOVD (Fokkema et al. 2011) locus-specific databases for APC and MSH2, and RB1 (Lohmann and Gallie 1993)). Unless otherwise medaled, variants predicted to be tolerated by *in silico* algorithms are assigned a bronze medal. Imperfect database matches (e.g., a different allele at the same genomic position or at the same codon but with a different amino acid change) are typically assigned a lower grade medal, e.g. silver rather than gold. Variants not meeting any of the previous criteria, e.g. most silent variants and those without any functional annotations, will not receive a medal. Amino acid and pathogenicity codes from the diverse variant databases used in this process are standardized to improve the reliability of annotations and utility of information (Methods). A summary of resources is shown in Table 1. MedalCeremony may also be run as a stand-alone pipeline on the St. Jude Cloud platform (Methods).

**Figure 2.**
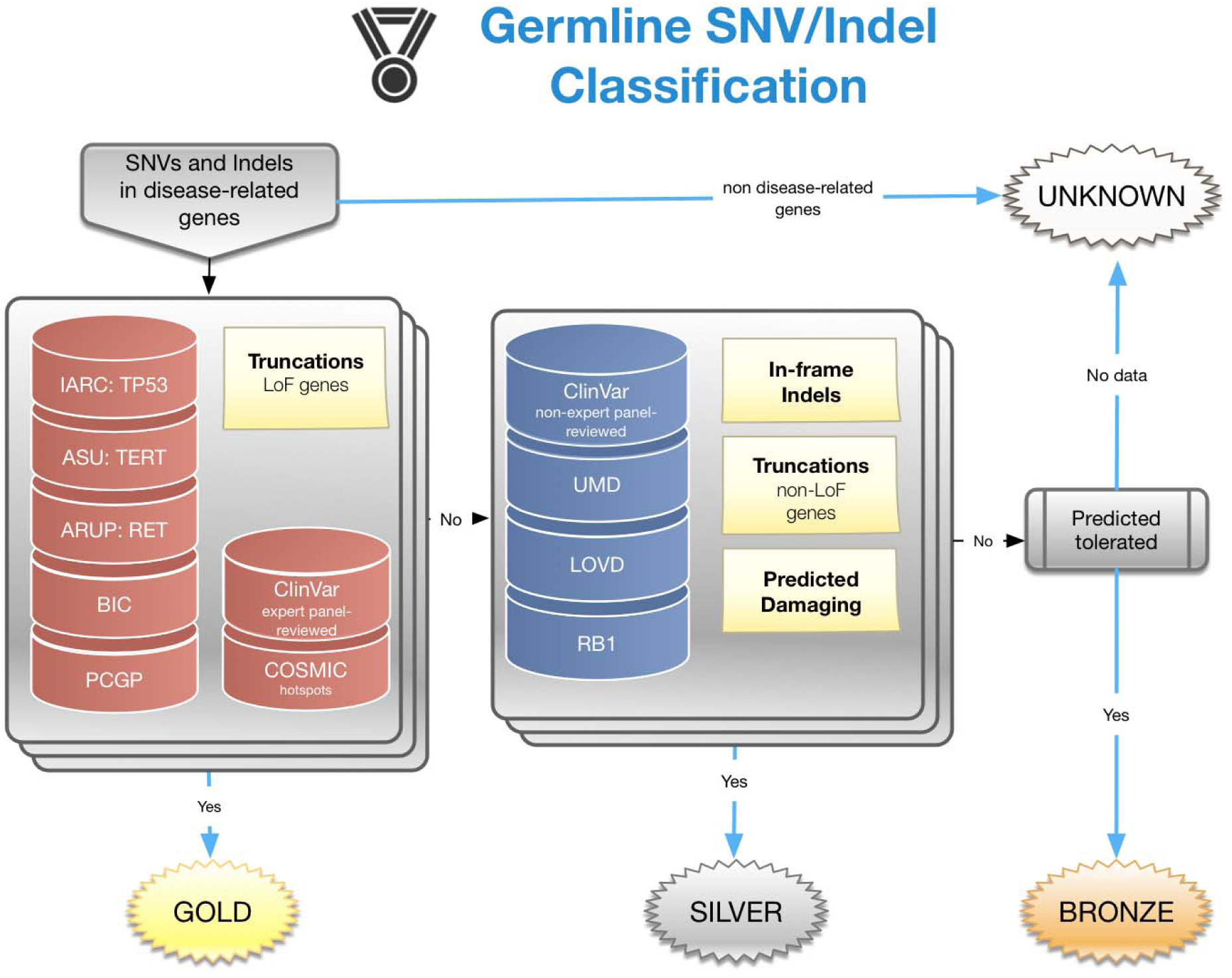
Design of the MedalCeremony pipeline for automated germline variant classification. Truncating variants in loss-of-function genes (e.g. tumor suppressors) and those matching highly-curated databases receive gold medals. Truncations in non-loss-of-function genes, in-frame indels, predicted damaging variants, and matches to additional databases receive silver medals. Otherwise variants predicted to be tolerated by damage-prediction algorithms receive bronze. Imperfect database matches receive a lower-grade medal than exact matches. Variants not meeting any of the prior criteria receive a result of “unknown”.

**Table 1.**
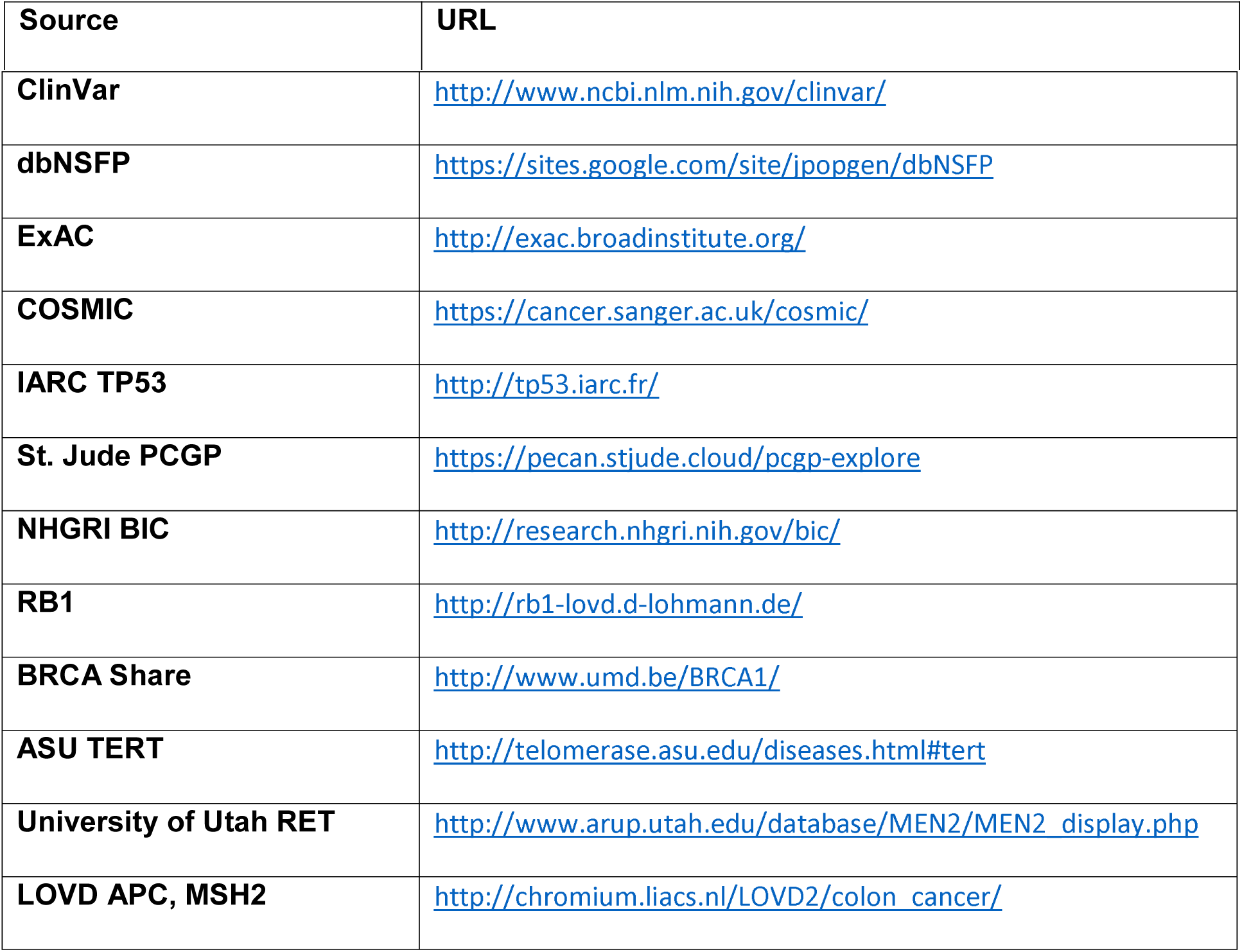
Databases used in classification

| Source | URL |
| --- | --- |
| **ClinVar** | http://www.ncbi.nlm.nih.gov/clinvar/ |
| **dbNSFP** | https://sites.google.com/site/jpopgen/dbNSFP |
| **ExAC** | http://exac.broadinstitute.org/ |
| **COSMIC** | https://cancer.sanger.ac.uk/cosmic/ |
| **IARC TP53** | http://tp53.iarc.fr/ |
| **St. Jude PCGP** | https://pecan.stjude.cloud/pcgp-explore |
| **NHGRI BIC** | http://research.nhgri.nih.gov/bic/ |
| **RB1** | http://rb1-lovd.d-lohmann.de/ |
| **BRCA Share** | http://www.umd.be/BRCA1/ |
| **ASU TERT** | http://telomerase.asu.edu/diseases.html#tert |
| **University of Utah RET** | http://www.arup.utah.edu/database/MEN2/MEN2_display.php |
| **LOVD APC, MSH2** | http://chromium.liacs.nl/LOVD2/colon_cancer/ |

### Variant review interface

After MedalCeremony classification, the results are presented in a table that can be searched or filtered by gene, variant class, medal status, or classification by expert review (Fig. 3A). If a variant has been previously classified by the user or the St. Jude germline variant review committee, that information will be pre-populated. Each row links to a variant page containing extensive annotations, including gene information from NCBI and OMIM (Amberger et al. 2015), ClinVar match details, population frequency, and *in silico* predictions of deleteriousness (Fig. 3B). The page also includes an embedded ProteinPaint view (Zhou et al. 2015), which overlays the current variant with aggregated somatic mutations and expert-classified P/LP germline variants on the protein product. This enables visual inspection of variant recurrence, hotspots, and enrichment of loss-of-function mutations.

**Figure 3.**
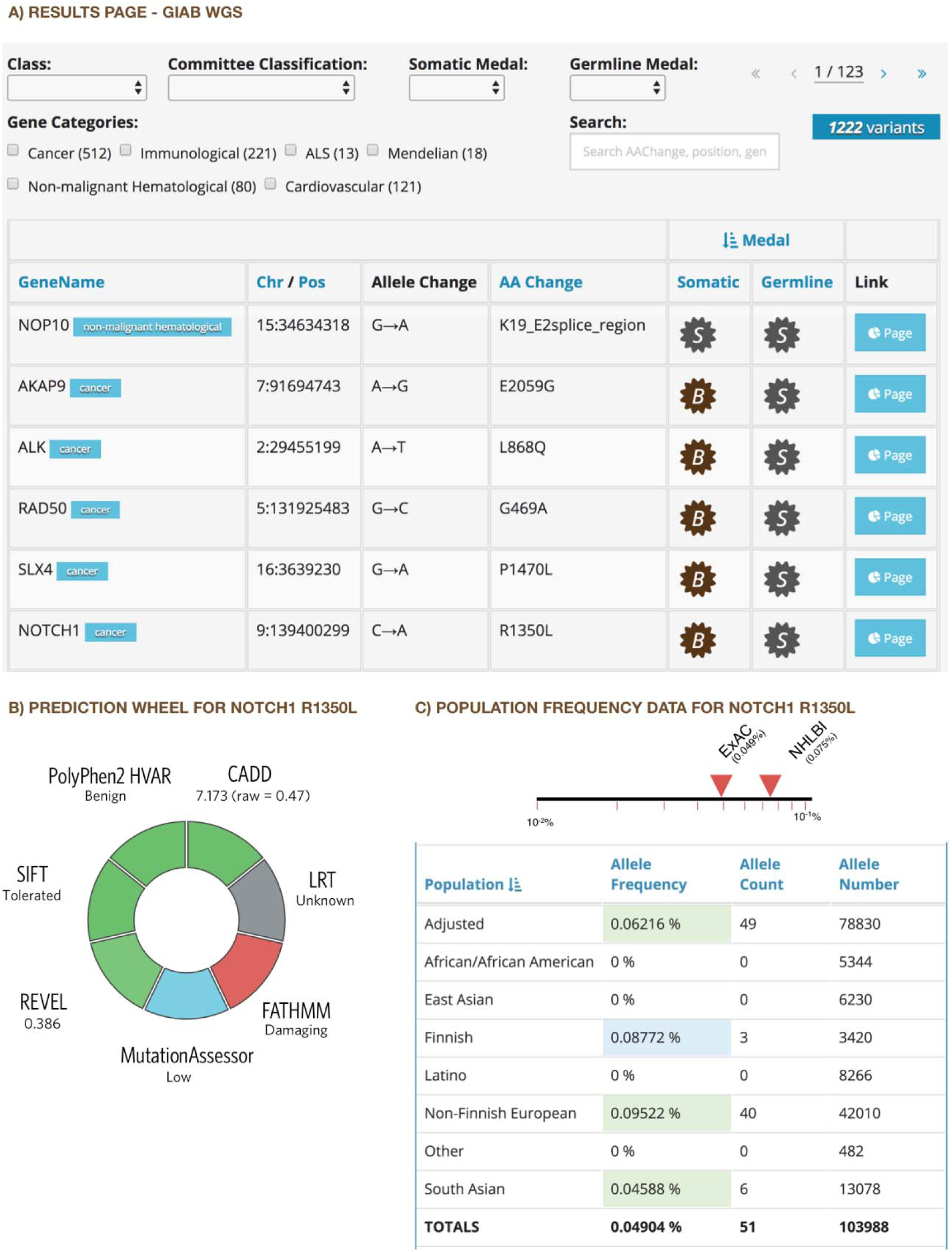
Annotation interface. Excerpts of PeCanPIE annotation interface. (A) Results for Genome in a Bottle WGS dataset. Variant page details for *NOTCH1* R1350L: (B) functional predictions, and (C) variant population frequency detail from ExAC ex-TCGA database.

**Figure 4.**
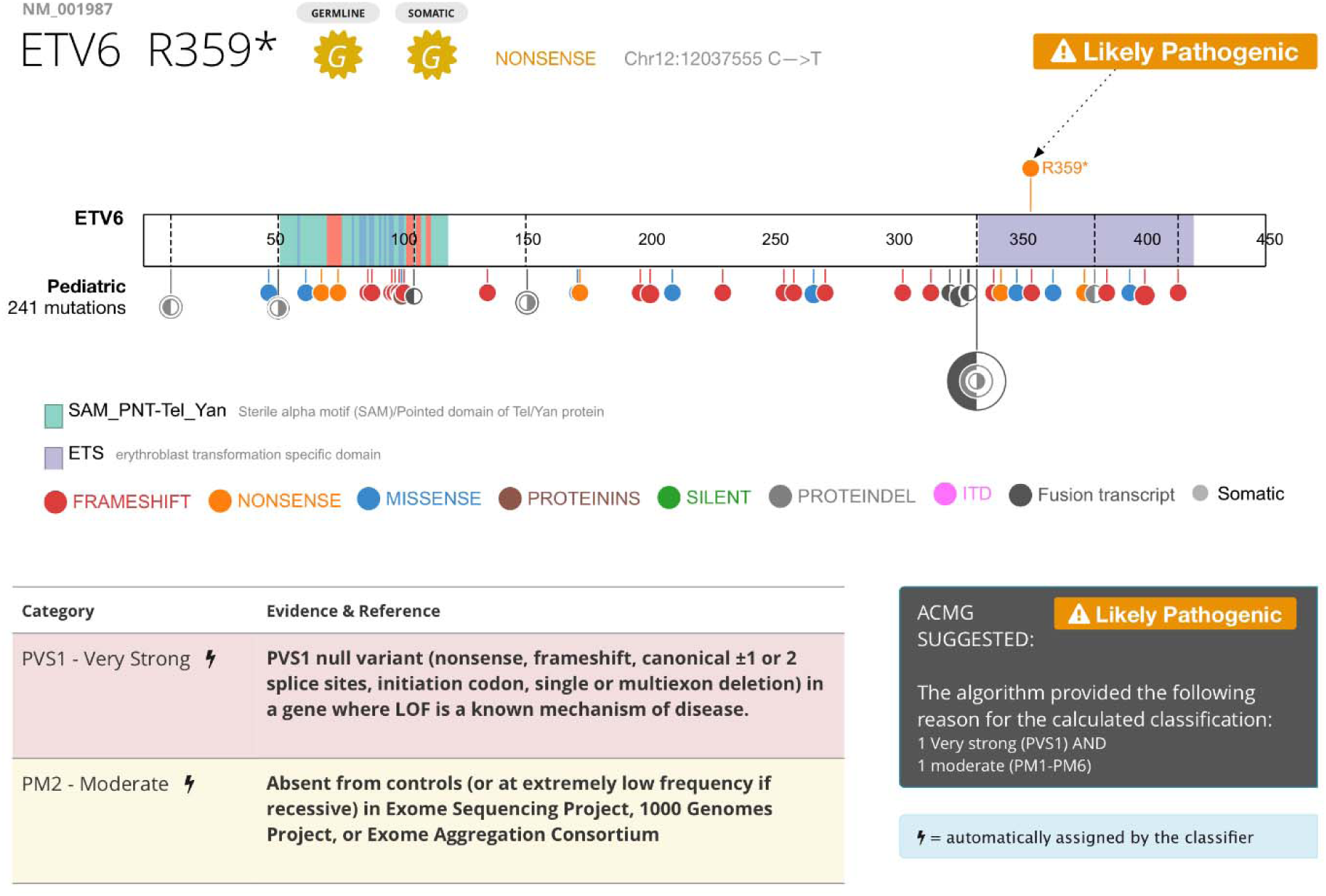
ACMG classification on ETV6. Top, ProteinPaint display of somatic *ETV6* variants across 11 subtypes of pediatric leukemia, showing enrichment of loss-of-function mutations (frameshifts in red, nonsense variants in orange). Arrow indicates position of germline R359* variant. Bottom, detail of PeCanPIE ACMG classification interface for R359* variant.

### ACMG classification interface

A powerful feature of the variant detail page is an interactive graphical interface that allows a reviewer to enter a series of pathogenicity criteria evidence tags (e.g., population frequency, segregation, functional significance, and *in silico* prediction), along with supporting information such as PubMed IDs, to automatically calculate a 5-tier classification: Pathogenic (P), Likely Pathogenic (LP), Unknown Significance (VUS), Likely Benign (LB), and Benign (B) based on the ACMG algorithm. MedalCeremony can automatically generate ACMG classification tags for variants, which are prepopulated into PeCanPIE’s classification interface. The following automatic tags are implemented: PVS1 (truncating variant in a tumor suppressor or other loss-of-function gene), PM1 (somatic hotspot in COSMIC), PM2 (absent from ExAC or appearing at a frequency of no greater than 0.0001) and the companion BA1 tag (>5% population frequency in ExAC), PM4 (in-frame protein insertions and deletions), PS1 and PM5 (amino acid comparisons made vs. pathogenic variants in ClinVar or those identified by the St. Jude Germline Review Committee). Automatically-assigned tags may be removed by the analyst if desired. This automation provides improved support versus manual curation interfaces, while still retaining analyst control over the ultimate classification decisions. As shown on the variant page for *ETV6* Arg359Ter, the single gold-medal variant detected in the patient with ALL was expert-classified as likely pathogenic because the mutation is present in a disease-related gene (i.e., *ETV6* is a pediatric ALL driver gene), is a loss-of-function null variant, and is not present in the ExAC database (Fig. 4).

Comparison of a germline variant with aggregated somatic variants can help inform germline classification for cancer predisposition genes. For example, family studies have identified a *PAX5* G183S germline mutation conferring susceptibility to B-ALL, which corresponds to somatic mutations detected in pediatric B-ALL and lymphoma (Shah et al. 2013). A similar profile was observed in the example WES data from an ALL patient presented in Fig. 1B: MedalCeremony assigned a single gold medal—a novel *ETV6* nonsense variant within the ETS domain (NM_001987.4:c.1075C>T, NP_001978.1:p.Arg359Ter)—based on the criteria of truncation in a tumor suppressor gene. The ProteinPaint view embedded in the variant page confirmed that in *ETV6,* somatic mutations are dominated by loss-of-function mutations across pediatric leukemia (Fig. 4), consistent with the tumor-suppressor gene model. Reviewers may enter custom evidence such as this into the interface for use during final classification.

### Pathogenicity classification of cancer predisposition genes in 4,000 pediatric cancer patients

PeCanPIE was designed in support of large-scale germline variation analysis projects, and was iteratively improved based on the feedback of an interdisciplinary group of researchers. Germline variants from the following studies have been analyzed thus far: 1) a study of germline variations in predisposition genes in 1,120 children with cancer (Zhang et al. 2015) classified 890 variants, identifying 109 as pathogenic (P) and 25 as likely-pathogenic (LP); 2) the St. Jude LIFE project, a follow-up study of 3,006 long-term survivors of pediatric cancer (Wang et al. 2018), classified 3,417 variants, including 188 P and 160 LP; and 3) Genomes for Kids (manuscript in preparation), a clinical research study of 310 pediatric cancer patients (https://clinicaltrials.gov/ct2/show/NCT02530658), clinically reported 25 P and 6 LP variants. PeCanPIE also serves as a repository for expert-curated decisions for the first two studies, whose resulting annotations are reapplied to incoming variant classification requests.

## Discussion

Although PeCanPIE’s features partially overlap those of other available tools (Li and Wang 2017; Masica et al. 2017), it provides several new capabilities. Specifically, variant classification is tightly integrated with the rich resource of somatic mutation data in pediatric cancer, which can be explored online via the embedded ProteinPaint view. Users can also analyze indels, MNVs, and complex substitutions, whereas web-based implementations of similar tools may be limited to SNVs alone (Li and Wang 2017). Another key feature is the cloud-based implementation of PeCanPIE, which obviates the need for complex software installation and command-line workflows. This design also allows back end analysis pipelines to be invoked independently from PeCanPIE, for users who prefer direct or programmatic access over a graphical interface. In comparison with web-based systems (Masica et al. 2017) which provide batch annotation of variants based on machine-learning scores (Carter et al. 2013, 2009), PeCanPIE provides more granular annotations and individual ACMG-recommended evidence tags to facilitate interpretation of pathogenicity classifications. Via dbNSFP, PeCanPIE also provides access to REVEL (Ioannidis et al. 2016) pathogenicity scores, which fared well in a recent comparison of algorithms for use with ACMG clinical variant interpretation guidelines (Ghosh et al. 2017). Lastly, PeCanPIE’s workflow offers advantages over CIVIC’s crowdsourced clinical interpretation of variants (Ta 2017), which relies on completely manual classification and data entry, i.e., VCF upload, annotation, and prioritization are not provided.

A limitation of the existing method is that damage-prediction algorithm scores are taken from the dbNSFP database, which only contains data for non-silent SNVs. While these annotations are unavailable for indels, because protein class annotations are taken into account by the scoring algorithm, high-impact events such as truncating variations will still be highly ranked. For variant population frequency filtering, we are currently using the TCGA-subtracted release of ExAC instead of gnomAD (Lek et al. 2016) because the gnomAD database contains TCGA samples; we plan to migrate to gnomAD once a TCGA-subtracted version becomes publicly available.

In conclusion, the PeCanPIE platform significantly accelerates the variant classification process by automating many prerequisite steps, helping to prioritize potentially pathogenic variants in NGS data, and providing a robust platform for investigating variant pathogenicity in disease-related genes. While PeCanPIE was developed and tested with pediatric cancer susceptibility as a primary focus, we are in the process of expanding its scope to other pediatric and adult diseases. Users are now able to specify custom gene lists to analyze appropriate to their diseases of interest, enabling disease-specific variant curation and facilitating gene discovery.

## Methods

### Disease-related gene list

The disease-related gene list comprises both cancer-related and non-cancer genes (Table S1). The cancer gene list was compiled from public resources and cancer genetic studies including: 1) studies of germline mutations in predisposition genes in cancer patients (Zhang et al. 2015; Huang et al. 2018; Wang et al. 2018); 2) cancer predisposition genes compiled by Rahman (Rahman 2014); 3) the Cancer Gene Census (Futreal et al. 2004); and 4) driver genes identified in pediatric and adult pan-cancer studies (Ma et al. 2018; Gröbner et al. 2018). Publications were reviewed to confirm the presence of either loss-of-function or gain-of-function mutations in cancer driver genes, excluding those previously identified as having elevated mutation rates (e.g. *LRP1B* (Lawrence et al. 2013)) and those reported only as fusion partners. Other disease-related genes include non-malignant hematological, immunodeficiency, and amyotrophic lateral sclerosis (ALS)-related genes (Taylor et al. 2016), and genes from ACMG and Ambry Genetics incidental finding gene lists (Kalia et al. 2017). Filtering the variants to disease-related genes helps focus on areas with relevant research interest and reduce the downstream processing burden, which is especially helpful for WGS data which may contain 4-5 million variants per sample. A user may choose to focus on one or more of these pre-defined disease categories for expert review or provide their own gene lists for custom analysis.

### Gene annotation and splice calling enhancement

Gene annotations are performed using the Ensembl Variant Effect Predictor (VEP) pipeline (McLaren et al. 2016), which provides information on a variant basis for the affected gene and transcript, functional class (e.g., silent, missense, and nonsense), and effect on protein coding. We enhanced splice variant annotation by reclassifying silent or missense variants at exon boundaries, which may impact splicing (e.g., *TP53* NM_000546.5:c.375G>A, NP_000537.3:p.Thr125Thr (Soudon et al. 1991)). While certainly not all of these variants will ultimately prove to be splice-related, these adjustments ensure additional scrutiny during expert review. A subsequent filtering step retains only variants in coding and splice-related regions. Silent variants are also kept because, in rare cases, they may cause aberrant splicing and thus be pathogenic. For example, ClinVar (Landrum et al. 2018) ID 90407 is a “silent” variant in the colon cancer predisposition gene *MLH1* (NM_000249.3:c.882C>T, NP_000240.1:p.Leu294=) that has been determined by an expert panel to be a pathogenic splice variant (Auclair et al. 2006). We refer to this enhanced pipeline as VEP+, which may also be run separately on the St. Jude Cloud platform.

### St. Jude Cloud platform

While PeCanPIE was designed as a web portal to maximize ease of use for non-bioinformaticians, two component pipelines are also publicly accessible. On its back end, St. Jude Cloud (https://stjude.cloud) uses DNAnexus (https://www.dnanexus.com/), a platform where user-created software pipelines can be installed and run on cloud computing instances. A DNAnexus account is required to use PeCanPIE for secure storage and to send notifications when submitted jobs are complete. Once a pipeline has been installed on DNAnexus, it is straightforward for non-expert users to run it, either from a standardized web interface or a command-line client. We have created two DNAnexus pipelines that are used by PeCanPIE, VEP+ for variant annotation (app-stjude_vep_plus) and MedalCeremony for automated classification (app-stjude_medal_ceremony). The availability of these component pipelines on the cloud provides users and institutions straightforward, scalable access to the software, and our centralized maintenance allows all users to immediately benefit from updates and new features as they become available. PeCanPIE is free for non-commercial use.

### Nomenclature standardization

We have observed that various variant databases which form the foundation of annotations for PeCanPIE vary in the structure and quality of variant specification. For example, databases may provide only protein-level annotations, only genomic annotations, or both. Likewise, there are many variations on the HGVS-like protein annotation nomenclature in circulation. The PeCanPIE code attempts to be flexible in parsing, standardizing, and formatting where possible, e.g. protein annotations may use either 3-character or 1-character protein codes (e.g. “Ser” or “S”), and a number of variations on stop codon formatting have been observed (“Ter”, “Term”, “*”, “X”, and “Stop”). In some cases partial information such as codon numbers were extracted from an otherwise incomplete annotation. Some databases also provide variations on the 5-tier ACMG pathogenicity calls which PeCanPIE attempts to standardize into B/LB/VUS/LP/P for easier comparison. We believe these standardizations further improve the reliability of annotations and utility of information provided by the PeCanPIE platform.

### Example data

The ALL variants in Figure 1b were called from St. Jude sample SJNORM015857_G1. Variant calling was performed with Bambino using the “high 20” profile which consists of the following command-line parameters: “-min-quality 20 -min-flanking-quality 20 -min-alt-allele-count 3 -min-minor-frequency 0 -broad-min-quality 10 -mmf-max-hq-mismatches 4 -mmf-max-hq-mismatches-xt-u 10 -mmf-min-quality 15 -mmf-max-any-mismatches 6 -unique-filter-coverage 2 -no-strand-skew-filter”. The results were subsequently filtered to variants having a variant allele frequency of at least 20%, an average mapping quality of 20 for variant reads, at least 5 reads of coverage for the variant allele, bi-directional confirmation of the variant allele, and at least 20 reads of total coverage. The results were converted to VCF by an in-house script and uploaded to PeCanPIE. The Genome-in-a-Bottle VCF used for Figure 1c is available from ftp://ftptrace.ncbi.nlmc/giab/ftp/release/NA12878_HG001/NISTv3.3.2/GRCh37/HG001_GRCh37_GIAB_highconf_CG-IllFB-IllGATKHC-Ion-10X-SOLID_CHROM1-X_v.3.3.2_highconf_PGandRTGphasetransfer.vcf.gz. This bgzip-compressed VCF file may be used directly with PeCanPIE.

### Software Availability

PeCanPIE is available at https://platform.stjude.cloud/tools/pecan_pie and is one component of the St. Jude Cloud platform (https://stjude.cloud/).

## Acknowledgements

This project was supported by the American Lebanese Syrian Associated Charities (ALSAC) of St. Jude Children’s Research Hospital, by a Cancer Center Support (Core) grant (CA21765) and a grant to JZ (CA21635) from the National Cancer Institute. We thank Yiyang Wu for discussions of *in silico* algorithms.

## Author contributions

Analysis pipeline design and development (M.N.E.), web software design and development (A.N.P., X.Z., J.B.B.), cloud pipeline development (M.N.E., C.L.M.), tool development (M.N.E., S.V.R., M.C.R.), genomic data analysis (E.R., D.J.H., Y.L., C.A.K., J.Z., S.N., Z.W., L.L.R., A.N.P., M.N.E.), manuscript text (M.N.E., J.Z., E.R., C.A.K.), figure preparation (A.N.P., M.N.E., J.Z.), database support (M.R.W.), project direction and supervision (J.Z., J.R.D., K.E.N.)

